# *C. elegans* granulins promote an age-associated decline in protein homeostasis via lysosomal protease inhibition

**DOI:** 10.1101/472258

**Authors:** Victoria J. Butler, Wilian A. Cortopassi, Andrea R. Argouarch, M. Olivia Pierce, Mihir Vohra, Juan A. Oses-Prieto, Fuying Gao, Benjamin Caballero, Shreya Chand, William W. Seeley, Bruce L. Miller, Giovanni Coppola, Alma L. Burlingame, Kaveh Ashrafi, Ana Maria Cuervo, Matthew P. Jacobson, Aimee W. Kao

## Abstract

The progressive failure of protein homeostasis is a hallmark of aging and a common feature in neurodegenerative disease. As the enzymes executing the final stages of autophagy, lysosomal proteases (or cathepsins) are key contributors to maintenance of protein homeostasis with age. Here, we identify the cysteine-rich granulin peptides as a new class of regulators of lysosomal aspartyl protease activity. Granulins are produced in an age and stress-dependent manner through cleavage of the neurodegenerative disease protein, progranulin. Once liberated, granulins selectively interact with the aspartyl protease ASP-3/cathepsin D to impair enzymatic activity. Consequently, protein homeostasis and lysosome function is disrupted, prompting cells to activate a compensatory transcriptional program. Our results support a model in which granulin production modulates a critical transition between the normal, physiological regulation of protease activity and the impairment of lysosomal function that can occur with age and disease.

## Introduction

Conventionally, aging and stress are thought to advance neurodegenerative disease risk through the accumulation of misfolded and aggregated proteins (David et al., 2010, Reis-Rodrigues et al., 2012, Walther et al., 2015), although understanding of the molecular basis of this phenomenon remains incomplete. Genetic and functional studies implicate age-related failures of protein degradation by the lysosome in the pathogenesis of multiple neurodegenerative diseases (Gotzl et al., 2016, Nixon, 2013, Wyss-Coray, 2016). Lysosomes contain fifteen cathepsins, which are defined by their active site nucleophile as serine, cysteine or aspartyl proteases (McDonald, 1986). These enzymes have a crucial role in processing and degrading proteins (Stoka et al., 2016). As their action is irreversible, protease activity needs to be tightly regulated. Serine and cysteine cathepsins undergo inhibitory regulation via steric association with small, cysteine-rich peptides known as serpins and cystatins, respectively (Silverman et al., 2001, Turk and Bode, 1991). To date, no such endogenous regulators of mammalian aspartyl proteases have been described.

Heterozygous progranulin (*PGRN*) loss-of-function mutations lead to autosomal dominant transmission of the neurodegenerative disorder frontotemporal lobar degeneration (FTLD) with TAR DNA binding protein 43 (TDP-43) inclusions (Neumann et al., 2006, Baker et al., 2006, Cruts et al., 2006). The molecular function of the progranulin protein remained elusive until it was indelibly linked to lysosomal function by the finding that loss of both gene alleles results in the lysosomal storage disease, neuronal ceroid lipofuscinosis (Smith et al., 2012). Progranulin localizes to lysosomes (Hu et al., 2010, Tanaka et al., 2013, Zhou et al., 2015, Gowrishankar et al., 2015) where it may act to promote lysosomal biogenesis and function (Tanaka et al., 2013, Tanaka et al., 2017, Evers et al., 2017, Beel et al., 2017). Furthermore, functional variants in progranulin have been implicated in lifespan regulation (Ahmed et al., 2010, Valenzano et al., 2015). The lentiviral delivery of progranulin to degenerating brain regions protects against neurotoxicity and cognitive defects in mouse models of Parkinson’s disease (Van Kampen et al., 2014) and Alzheimer’s disease (Minami et al., 2014). As such, efforts to increase progranulin production in patients are underway (Alberici et al., 2014, Cenik et al., 2011, Sha et al., 2017, She et al., 2017).

The progranulin (PGRN) protein can be proteolytically cleaved to liberate multiple small, cysteine-rich “granulin” peptides (Plowman et al., 1992). Granulins are highly conserved, disulfide-bonded miniproteins with unknown biological function. (Kao et al., 2017, Lavergne et al., 2012, Palfree et al., 2015, Cenik et al., 2012, Bateman and Bennett, 2009). Owing to the twelve cysteines and six disulfide bonds found in each cleaved granulin, these peptides adopt a stacked β-sheet configuration that is compact, structurally stable and potentially protease resistant (Tolkatchev et al., 2008). Several lines of evidence exist that cleaved granulin peptides oppose the function of the full-length protein. While progranulin has proliferative (He and Bateman, 2003, Tolkatchev et al., 2008) and anti-inflammatory (Kessenbrock et al., 2008, Zhu et al., 2002) properties, granulin peptides have been shown to inhibit cell growth (Tolkatchev et al., 2008) and stimulate inflammation (Zhu et al., 2002). In addition, we have previously demonstrated a role for granulins in selectively promoting the accumulation of TDP-43, thereby exacerbating TDP-43 toxicity and potentially contributing to the pathogenesis of disease (Salazar et al., 2015). However, the mechanism by which granulins exert this specific regulation on TDP-43 metabolism is completely unknown.

Here, we show that granulins regulate TDP-43 degradation and endolysosomal function via the selective impairment of the lysosomal aspartyl protease ASP-3/cathepsin D (CTSD). Although granulin function, like that of cystatins and serpins, is required for normal development (Stoka et al., 2016), increased production with age and stress reduces animal fitness by disrupting protein homeostasis. Overall, our findings highlight granulins as critical regulators of proteolytic lysosomal function and potential drivers of neurodegenerative disease pathogenesis.

## Results

### Granulins impair organismal fitness and are produced in an age- and stress-dependent fashion

We have recently shown that *C. elegans* progranulin *(pgrn-1)* null mutants exhibit enhanced resistance to endoplasmic reticulum (ER) unfolded protein stress (Judy et al., 2013). As a genetic null, *pgrn-1(-)* animals produce neither full-length progranulin nor cleaved granulins; therefore, absence of either the holoprotein or the cleavage fragments could be responsible for the ER stress resistance. Based on our earlier finding that granulins could exacerbate TDP-43 toxicity (Salazar et al., 2015), we hypothesized that the bioactive granulins were responsible for inhibiting ER stress resistance. Hence, to isolate granulin activity, we expressed individual *C. elegans* granulins 1, 2 and 3 in a *pgrn-1* null background. Granulin expression in a progranulin null background completely abolished the ER stress resistance phenotype (Figure 1A). In contrast, animals over-expressing *C. elegans* full-length progranulin in a progranulin null background remained ER stress resistant (Figure 1B). This suggests that it is the granulins, and not full-length progranulin, that normally inhibit ER stress resistance.

**Figure 1.**
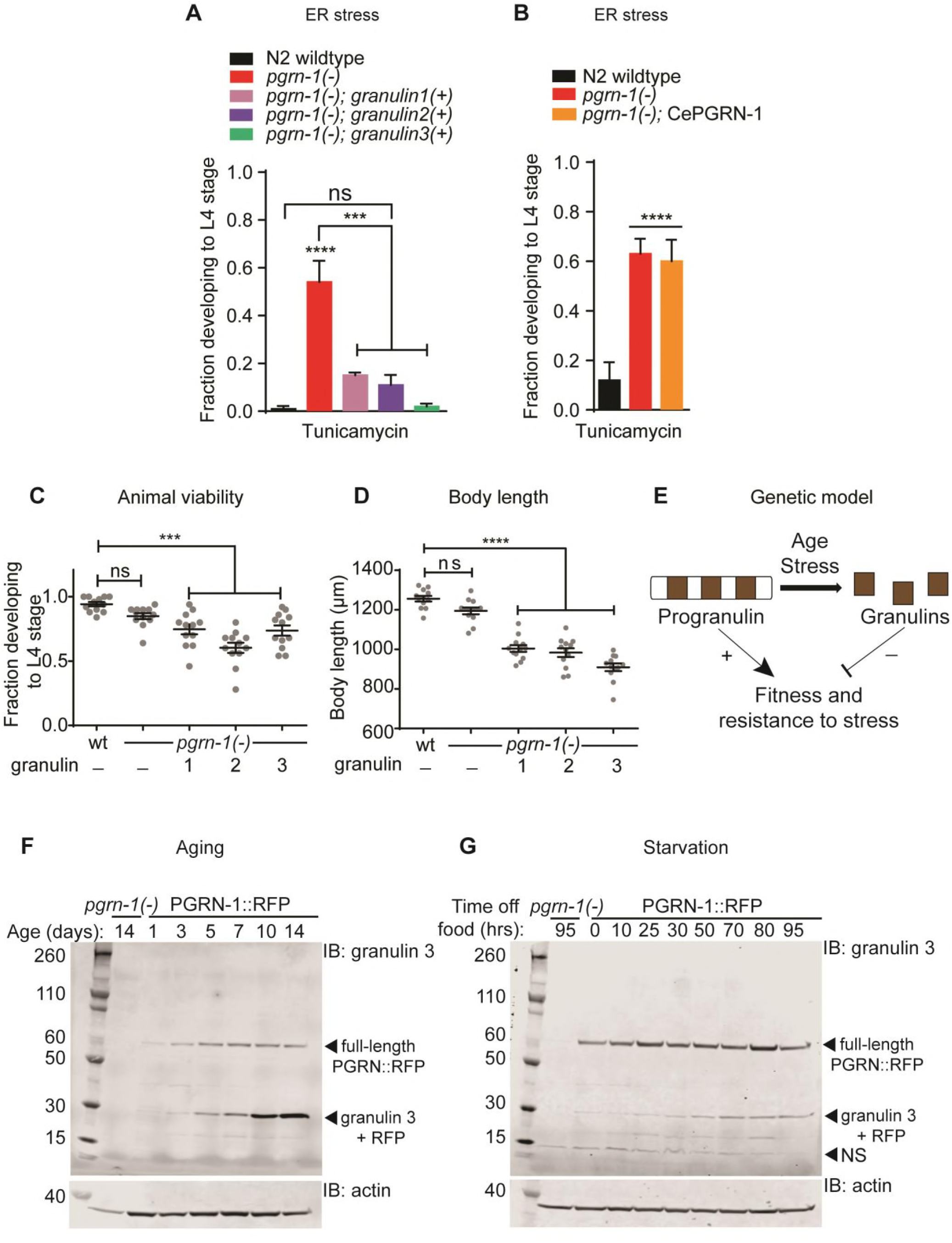
Granulins impair fitness and are produced in an age- and stress-dependent manner. (**A**) Wild-type (N2) and *pgrn-1(-)* animals with and without granulin expression were subjected to ER stress with tunicamycin (5 μg / ml). The fraction developing to L4 stage was quantified (n = 50, 3 biological replicates). (**B**) Wild-type (N2) and *pgrn-1(-)* animals with and without *C. elegans* progranulin over-expression were subjected to ER stress with tunicamycin (5 μg / ml). The fraction developing to L4 stage was quantified (n = 50, 3 biological replicates). (**C**) Wild-type and *pgrn-1(-)* animals with and without granulin expression were staged as embryos. Animals were scored for development to L4 stage (n = 50, 12 biological replicates). (**D**) Measurement of body length at day 1 adulthood (n = 12). (**E**) Genetic model for progranulin and granulin function in stress resistance. (**F-G**) Western blot of *C. elegans* PGRN-1::RFP lysates with aging (**F**), and starvation (**G**). Immunoblotting was performed with an anti-granulin 3 antibody. The full length PGRN::RFP at ~60kDa was recognized by antibodies against granulins 1, 2, 3, and RFP. NS = non-specific band. The most prominent cleavage product at ~30kDa was recognized by both granulin 3 and RFP antibodies (see Figure S1E). Throughout, bars show mean ± s.e.m., one or two-way ANOVA with post-hoc Tukey multiple comparisons test. Comparisons are to wild-type unless otherwise indicated (***P<0.001, ****P<0.0001, ns = not significant, wt = wild-type).

While working with the granulin-expressing lines, we noted a decrease in overall animal fitness attributable to the granulins. Granulin production significantly reduced animal viability by lowering the number of eggs that hatched and slowing the development of animals to maturity (Figure 1C). Granulin-expressing animals that did reach adulthood were smaller in size (Figure 1D). These data, coupled with previous work by others on the function of progranulin (He and Bateman, 2003, Kessenbrock et al., 2008, Tolkatchev et al., 2008, Zhu et al., 2002), suggest that granulins impair animal fitness and resistance to stress while progranulin promotes these qualities (Figure 1E).

In *C. elegans* and mammals, progranulin production increases with age (Kao et al., 2011, Petkau et al., 2010) and injury (Petkau and Leavitt, 2014, Holler et al., 2016). However, the degree to which granulin peptides are liberated has not been measured. We first asked if progranulin cleavage into granulins increases with age. Using a translational reporter, PGRN-1::RFP, we found that granulin production does indeed increase in an age-dependent fashion (Figures 1F, S1A and S1B). Granulin cleavage also increased in response to certain physiological stressors such as starvation (Figures 1G, S1C and S1D). Thus, age and stressful stimuli appear to promote the cleavage of full-length progranulin into granulins, which impair overall animal fitness.

### Granulins localize to the endolysosomal compartment

To establish the trafficking and localization of granulin peptides within a whole organism, we utilized microscopy and biochemistry techniques. First, we determined the sub-cellular localization of full-length progranulin using our translational progranulin reporter and organelle-specific markers. As expected, in cells that secrete progranulin, such as the intestine, the reporter co-localized with both a Golgi marker, mannosidase II (Figure 2A), and a lysosome marker, lysosomal-associated membrane protein 1 (LMP-1) (Figure 2B). However, in coelomocytes, a cell type that takes up but does not produce progranulin, the progranulin reporter was only seen in the endolysosomal compartment (Figures 2C-E), suggesting that extracellular progranulin is transported through endosomes to reach the lysosome.

**Figure 2.**
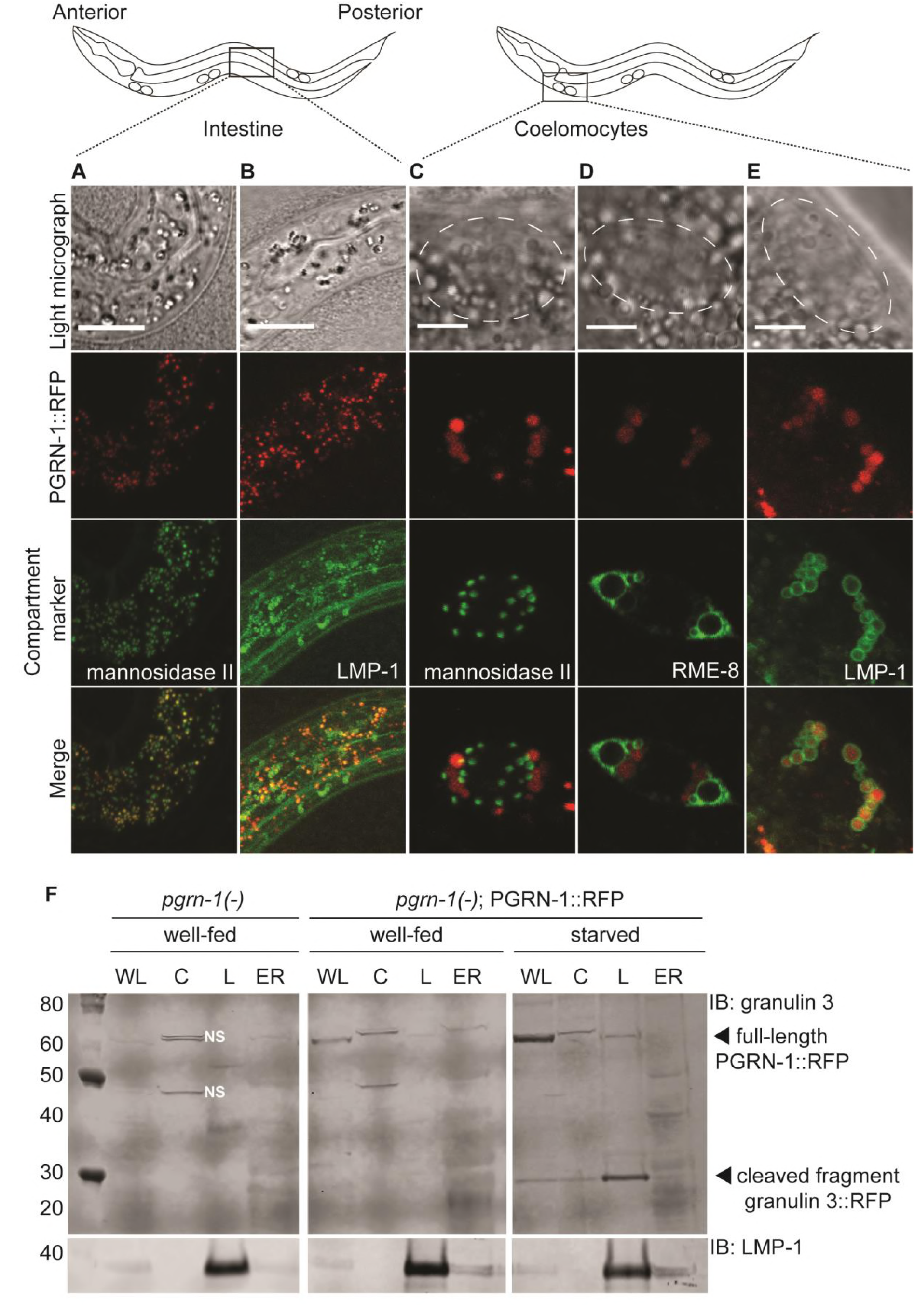
*C. elegans* progranulin and granulins localize to the endolysosomal compartment. A translational PGRN-1::RFP reporter (red) co-localizes with (**A**) Golgi (mannosidase II::GFP) and (**B**) lysosomes (LMP-1::GFP) in the intestine of L1 stage larvae (anterior to the left). In scavenging coelomocytes, PGRN-1::RFP does not co-localize with (**C**) Golgi (mannosidase II::GFP), but does co-localize with (**D**) early, late and recycling endosomes (RME-8::GFP) and (**E**) lysosomes (LMP-1::GFP). Dashed white lines mark the outline of each coelomocyte cell, scale bar = 5 μm. Shown are representative images from confocal microscopy z-stack sections taken at 0.7 μm. (**F**) Subcellular fractionation of PGRN::RFP animals. Whole lysate (WL), cytosol (C), lysosome (L) and endoplasmic reticulum (ER) fractions from fed and starved animals were immunoblotted with anti-granulin 3 and anti-LMP-1 antibodies. The same progranulin full-length and cleavage bands were also identified with an anti-RFP antibody (data not shown). Fed *pgrn-1(-)* animals are shown as a control for non-specific bands (NS).

Having established that progranulin can be trafficked from one tissue type to another, we next sought to better understand the subcellular localization of cleaved granulin peptides. To do so, we developed a protocol for subcellular fractionation of *C. elegans*. The purity of cytosolic, ER and endolysosomal fractions was confirmed with established markers (Figures S2A-C). Fractionation of fed or starved animals expressing the PGRN-1::RFP reporter was then performed. In fed animals, full-length progranulin was enriched in the endolysosomal fraction with very little lower molecular weight granulin observed in any fraction (Figure 2F). Upon starvation, the cleaved granulins increased primarily in the endolysosomal fraction, confirming that the majority of the age and stress-induced granulins are, in fact, endolysosomal (Figure 2F). Individual granulins that were transgenically expressed also demonstrated lysosomal localization (Figures S2D-F). Therefore, granulin peptides are produced *in vivo* in the endolysosomal compartment in a stress-responsive manner.

### Granulins impair protein homeostasis and activate the lysosomal CLEAR response

Given that granulins impair organismal fitness, localize to the endolysosomal fraction and impair stress resistance, we next investigated their impact on protein homeostasis and lysosomal function. First, we asked if granulin expression affects lysosomal biogenesis and morphology. In *C. elegans*, coelomocytes scavenge and detoxify the pseudocoelomic cavity and therefore have a well-developed endo-lysosomal system (Fares and Greenwald, 2001). Indeed, expression of granulin 3 grossly deformed these organelles (Figure 3A-D). Lysosomes lost their spherical shape and exhibited abnormal membrane protrusions and tubular extensions (Figure 3A-C), and became smaller and less numerous (Figure 3D).

**Figure 3.**
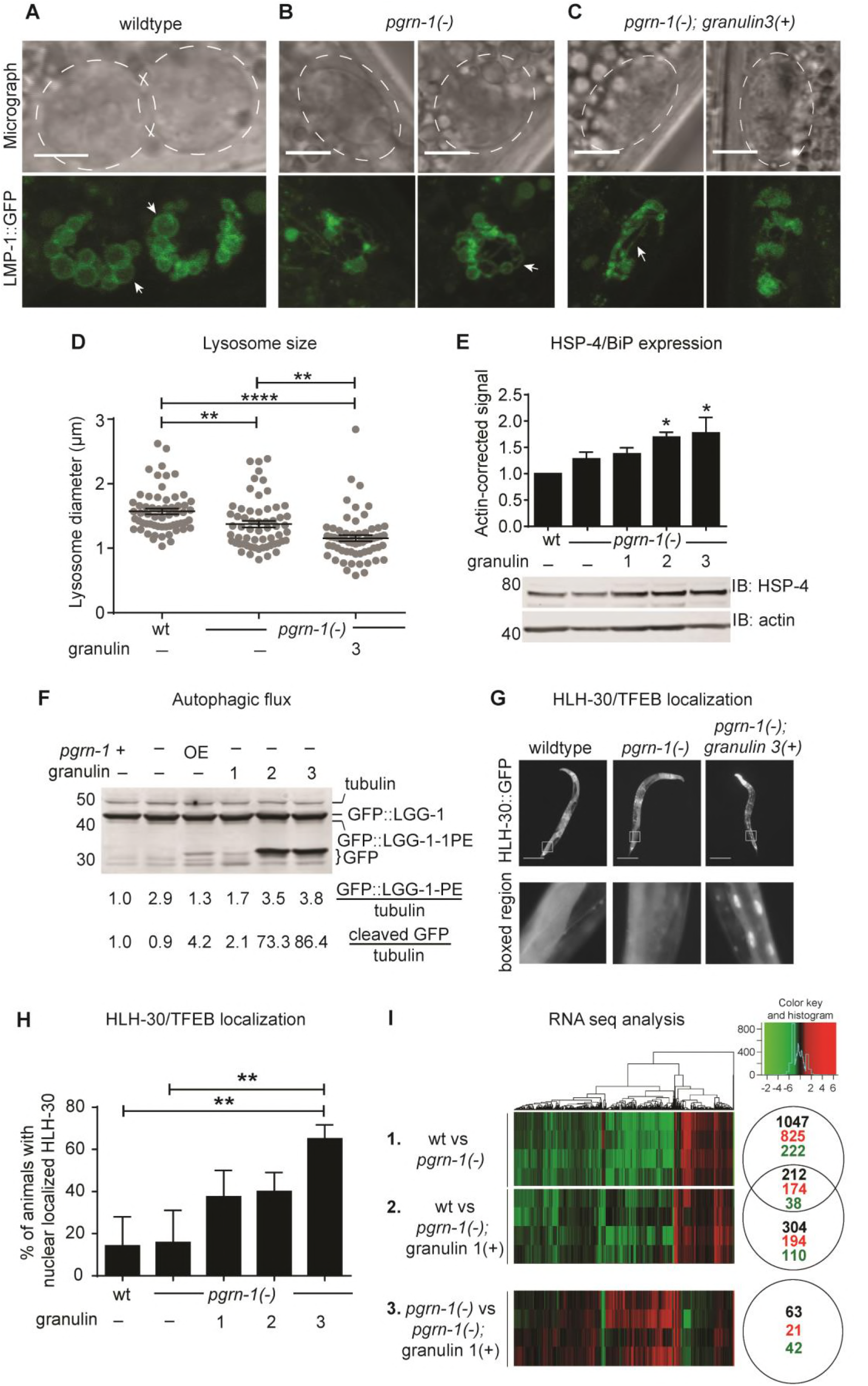
Cleaved granulins localize to the endo-lysosomal fraction and activate the CLEAR program. (**A-C**) Representative confocal images of anterior coelomocyte cells expressing LMP-1::GFP in (**A**) wild-type animals, (**B**), *pgrn-1(-)* animals and (**C**) *pgrn-1(-); granulin 3(+)* animals. Scale bars are 10 μm in the wild-type panel and 5 μm in the *pgrn-1(-)* and *pgrn-1(-); granulin 3(+)* panels. Dashed white lines mark the outline of each coelomocyte cell. White arrow heads indicate spherical lysosomes or abnormal tubular extensions. (**D**) Lysosome diameter measurements from anterior coelomocyte cells (n = 60). (**E**) Total worm lysates from synchronized day 1 adult granulin-expressing animals were immunoblotted with an anti-HSP-4/BiP antibody (3 biological replicates). Anti-actin was used as a loading control. (**F**) Total worm lysates from synchronized day 1 adult animals expressing GFP::LGG-1 were immunoblotted with an anti-GFP antibody. Anti-tubulin was used as a loading control. OE = animals over-expressing *C. elegans* full-length progranulin in a null background. Values shown are representative of three independent experiments. (**G**) Representative images of wild-type, *pgrn-1(-)* and *pgrn-1(-); granulin 3(+)* animals expressing HLH-30::GFP (Scale bar = 200 μm.) (**H**) Percentage of animals with nuclear localized HLH-30::GFP (n = 120 animals from 3 biological replicates). (**I**) Heat map showing the fold-changes of gene expression in comparisons of day 1 adult animals as indicated. Data from four independent biological replicates are shown. Significance cut-off was an FDR of <0.05. The numbers in black represent the total number of differentially expressed genes, with direction of change (up-regulated = red, down-regulated = green) indicated below. See **Supplementary Table 1** for the complete gene list.

Given that granulins impair lysosomal morphology, we wondered if they more broadly impacted protein homeostasis. Thus, we measured endogenous levels of heat shock protein HSP-4, the nematode BiP/Grp78 homolog (Heschl and Baillie, 1989). HSP-4/BiP expression is upregulated during the unfolded protein response (UPR) (Calfon et al., 2002). We found that granulin-expressing animals displayed a trend for increased basal expression of HSP-4/BiP on day 1 of adulthood, reaching significance in animals expressing granulin 2 and 3 (Figure 3E). Therefore, granulin expression perturbs protein homeostasis, induces the UPR and upregulates HSP-4 expression.

Next, we assessed autophagic flux by performing an immunoblotting analysis using a GFP-tagged LGG-1 protein (GFP::LGG-1) expressed under the endogenous *lgg-1* promoter (Kang et al., 2007, Melendez et al., 2003). LGG-1 is the Atg8/LC3 II homolog in *C. elegans* (Melendez et al., 2003). On conjugation to phosphatidylethanolamine (GFP-LGG-1::1-PE) the GFP::LGG-1-PE becomes associated with the autophagosome membrane. After fusion of the autophagosome with the lysosome, GFP::LGG-1-PE is degraded. However, the GFP fragment is relatively stable to proteolysis and accumulates in an autophagy-dependent manner (Gutierrez et al., 2007, Hosokawa et al., 2006, Shintani and Klionsky, 2004). Therefore, an increase in GFP::LGG-1-PE and cleaved GFP forms reflect an increase in autophagic flux (Klionsky et al., 2008). Immunoblotting demonstrated that both GFP::LGG-1-PE and cleaved GFP levels were increased in granulin 2 and 3 expressing animals. The increase in cleaved GFP was particularly striking, being over 70 times more than controls (Figure 3F). This strong enrichment of markers suggests an upregulation of autophagy in granulin 2 and 3 expressing animals. Interestingly, we previously observed that granulin 2 and 3 expression specifically enhanced TDP-43 toxicity, despite being expressed at similar levels to granulin 1 (Salazar et al., 2015).

Lysosomal biogenesis and autophagy are both regulated by the master lysosomal transcription factor, TFEB (Sardiello et al., 2009, Settembre et al., 2011). The *C. elegans* TFEB is HLH-30 (Lapierre et al., 2013). In response to starvation, stressful stimuli and aging, TFEB/HLH-30 translocates from the cytosol to the nucleus to activate its transcriptional targets (Sardiello et al., 2009, Settembre et al., 2011, Settembre et al., 2012, Lapierre et al., 2013). This program, known as the Coordinated Lysosomal Expression and Regulation (CLEAR) response induces expression of genes involved in lysosome function and autophagy, including progranulin. We observed HLH-30/TFEB localization in control, *pgrn-1(-)* and granulin expressing animals. Granulin expression promoted nuclear localization of TFEB/HLH-30, reaching significance in granulin 3-expressing animals (Figures 3G-H). This effect was not seen in *pgrn-1(-)* animals. These results suggest that the disruption of lysosome morphology and protein homeostasis seen in granulin-expressing animals leads to compensatory translocation of HLH-30/TFEB from the cytosol to the nucleus.

To determine if the granulin-induced TFEB/HLH-30 translocation also induced a transcriptional CLEAR program, we performed RNA seq profiling of wild-type, *pgrn-1(-)* and granulin-expressing animals (**Supplementary Table 1**). We first compared *pgrn-1(-) and pgrn-1(-); granulin* animals with wild-type animals. Loss of progranulin resulted in 1047 differentially expressed genes and granulin expression resulted in 304 differentially expressed genes, many of which were shared between the two strains (Figures 3I-1 and 3I-2). In contrast, when we compared *pgrn-1(-)* null animals with and without granulin expression, a distinct set of differentially regulated transcripts were observed (Figure 3I-3). The animals expressing granulin significantly upregulated genes whose promoters contained the putative TFEB/HLH-30 binding site E-box sequence (p = 0.009) (Grove et al., 2009, Lapierre et al., 2013) (**Supplementary Table 2**). GO term analysis showed enrichment in genes associated with growth rate and cell cycling (**Supplementary Table 1**). To determine if the upregulation of TFEB target genes was a compensatory transcriptional response in granulin-expressing animals, we crossed these animals into an *hlh-30(-)* null background. When lacking *hlh-30*, we observed that fewer granulin-expressing animals grew to adulthood and this was due to an increase in the number of animals arresting at early larval stages (Figure S3). Together, these data demonstrate that granulin expression, even in the absence of stress or starvation, was sufficient to activate a compensatory CLEAR response and induce expression of genes containing TFEB binding sites. Overall, the ability of granulins to 1) direct TFEB to the nucleus, 2) induce a CLEAR response and 3) disrupt lysosome morphology, indicates that granulin-dependent impairment of lysosomal function negatively impacts cellular protein homeostasis.

### Granulins interact with the aspartyl protease ASP-3/cathepsin D and impair its protease activity

To better understand the mechanism by which granulins exert their effect on lysosome function and protein homeostasis, we performed unbiased co-immunoprecipitation/mass spectrometry experiments using *C. elegans* granulin 3 as bait. The list of statistically enriched granulin 3 interacting proteins included several lysosomal proteins, including ASP-3, the nematode ortholog of mammalian cathepsin D (CTSD). (Supplementary Table 3). ASP-3/CTSD was of particular interest because it is a stress-activated aspartyl protease with allelic variants that increase risk for neurodegenerative disease (Paz et al., 2015, Schuur et al., 2011, Johansson et al., 2003). In order to gain insight into how granulins might interact with ASP-3/CTSD, we performed unbiased *in silico* computational modeling. Homology modeling of *C. elegans* granulins 1, 2, and 3, based on human granulin A and carp granulin 1 (Tolkatchev et al., 2000, Vranken et al., 1999), show that all three nematode proteins display the well-defined and evolutionarily-conserved stacked beta hairpin conformation of granulins (Figures 4A-C) (Tolkatchev et al., 2008). Protein-protein docking algorithms predicted that the most energetically favorable sites of interaction between each granulin and ASP-3/CTSD were through the propeptide region (Figure 4D-F). These data suggest that the granulin domains may interact with the propeptide of ASP-3/CTSD to regulate protease maturation or activity.

**Figure 4.**
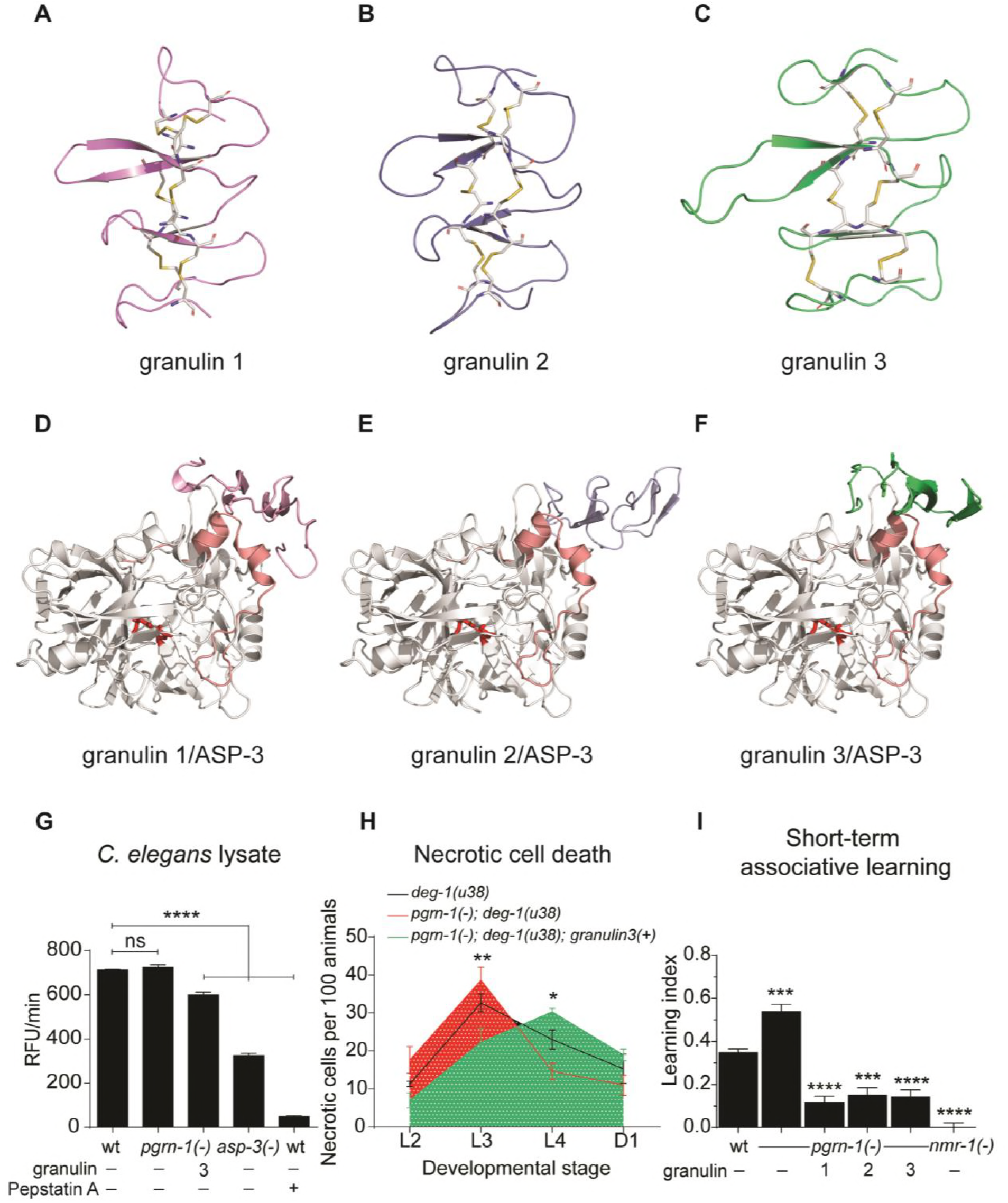
Granulin peptides impair cathepsin D activity. (**A-C**) Homology models for *C. elegans* granulin 1 (**A**), granulin 2 (**B**) and granulin 3 (**C**). Disulfide bonds forming the beta stacks are shown in yellow. (**D-F**) Non-covalent protein-protein docking results between *C. elegans* pro-ASP-3 (residues 23 to 392, in white) and granulins 1 (**D**), 2 (**E**) and 3 (**F**). Aspartate active site residues are shown in red and the propeptide in pink. (**G**) ASP-3/CTSD activity measured in total worm lysates (data from three technical replicates are shown, data is representative of three biological replicates, values shown are mean ± s.e.m., one-way ANOVA with Tukey multiple comparisons test, ****P<0.0001). Animals lacking *asp-3* continue to have significant activity, likely due to other aspartyl proteases that are inhibited by Pepstatin A. (**H**) The *deg-1(u38)-induced* necrosis of the mechanosensory PLM neurons is shifted to the right in granulin 3 expressing animals (n = 300, data pooled from 3 independent experiments.) Values shown are mean ± s.e.m., two-way ANOVA with Tukey multiple comparisons test, *P<0.05, **P<0.01, comparisons are to *pgrn-1(-); deg-1(u38))*. (**I**) Measurement of short-term associative learning (three biological replicates). The glutamate receptor mutant *nmr-1(ak4)* was used as a positive control. Values shown are mean ± s.e.m., one-way ANOVA with Tukey multiple comparisons test, ***P<0.001, ****P<0.0001.

Having established a physical interaction between granulins and ASP-3/CTSD, we next asked whether ASP-3/CTSD activity is changed by this interaction. To do so, we directly measured ASP-3/CTSD activity in lysates from granulin-expressing *C. elegans*. Granulin 3 expression significantly reduced ASP-3/CTSD protease activity (Figure 4G). ASP-3/CTSD is required for several biological processes in *C. elegans*, including necrotic cell death (Syntichaki et al., 2002) and for learning and memory in humans (Payton et al., 2003, Payton et al., 2006). Consistent with an inhibitory effect, granulin 3 delayed ASP-3/CTSD-dependent necrotic cell death (Figure 4H). Short-term associative learning can be assayed in *C. elegans* using a positive olfactory learning paradigm (Kauffman et al., 2010, Torayama et al., 2007). When granulin-producing animals were tested in this assay they underperformed compared to controls (Figure 4I), suggesting that granulin expression results in neuronal dysfunction. Recently, several studies have reported an interaction between full-length progranulin and CTSD (Beel et al., 2017, Valdez et al., 2017, Zhou et al., 2017a). Overall, our data suggest that granulin peptides can also interact with CTSD and have a functional inhibitory effect on aspartyl proteases *in vivo*.

### Patients with FTLD due to *Pgrn* mutations exhibit increased granulin fragments and decreased CTSD activity in affected brain regions

Thus far, our data demonstrate that excess granulins impair protein homeostasis via inhibition of aspartyl protease activity. To determine if granulin production contributes to neuronal dysfunction in FTLD due to *PGRN* mutations, we examined progranulin cleavage fragments and CTSD activity in post-mortem brain tissue from patients with FTLD due to progranulin mutations and controls (characteristics of subjects and matched controls are found in Table 1). As internal controls for each subject, we sampled from both a non-diseased and diseased brain region [inferior occipital cortex (IOC), gliosis score of 0-1 and middle frontal gyrus (MFG), gliosis score of 3, respectively]. We have previously shown that levels of a 33 kDa granulin fragment are higher in degenerating regions from patients with AD and FTLD (not due to *PGRN* mutations) (Salazar et al., 2015). Here, we similarly found that despite progranulin heterozygosity, the FTLD-TDP-*PGRN* subjects produced more of the 33 kDa granulin fragment in degenerating brain regions and overall more than control subjects (Figure 5A-B), likely due to copious progranulin production by infiltrating inflammatory cells. Immunoblotting for CTSD expression levels showed that antibodies against CTSD recognize both an immature pro-CTSD at 48kDa and a cleaved and mature CTSD at 30kDa. Both pro-CTSD and mature CTSD expression levels were significantly increased in degenerating brain regions and overall more than control subjects (Figure 5C-D). Despite the presence of more CTSD protein, CTSD activity in diseased-affected brain regions was *lower* in patients with FTLD-TDP-*PGRN* compared to controls (Figure 5E). These data show that FTLD-TDP-*PGRN* mutation carriers exhibit decreased CTSD activity in a diseased area of brain, and this correlates with increased progranulin cleavage fragments in this area.

**Figure 5.**
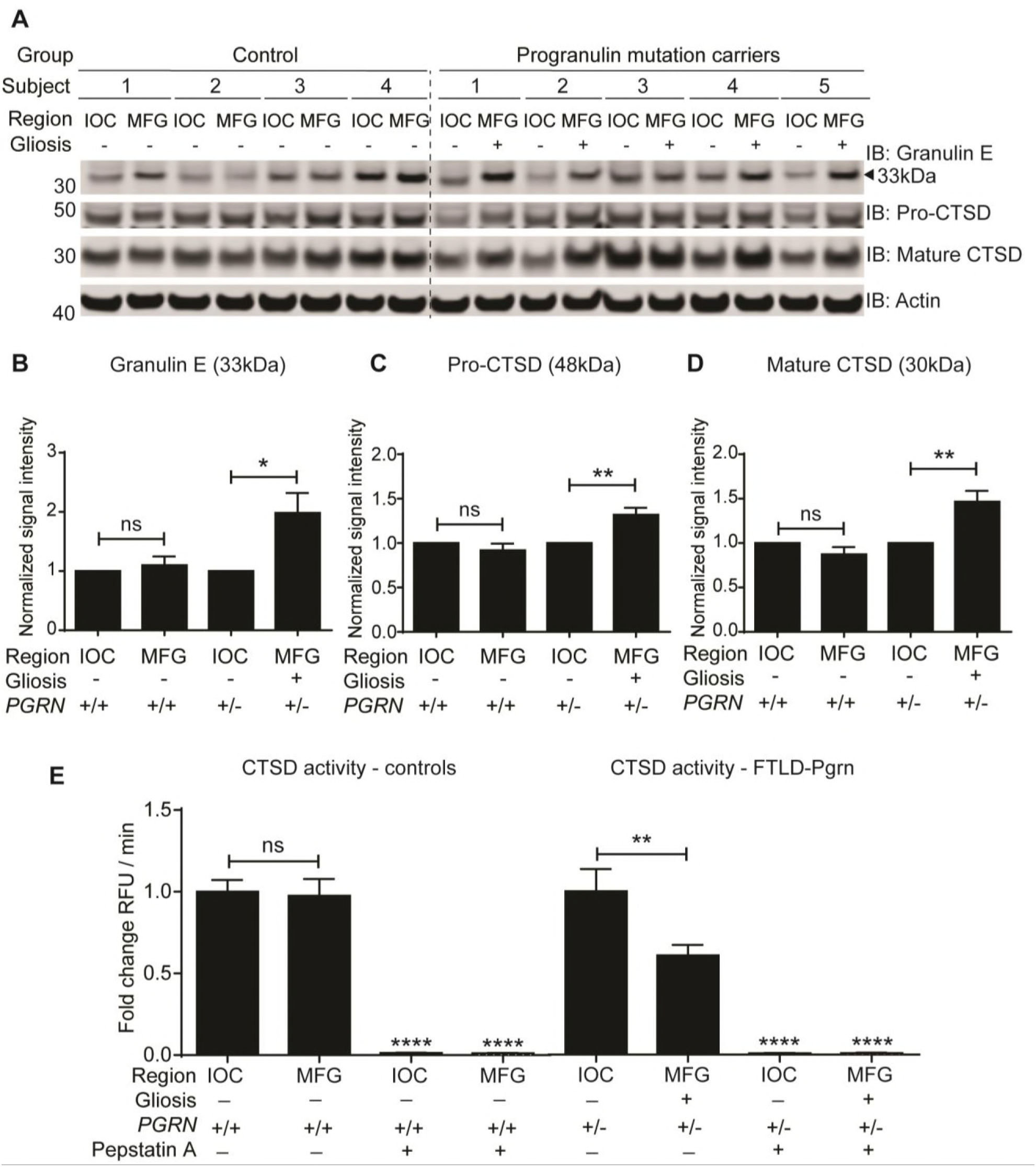
A cleaved granulin peptide is enriched in highly degenerative cortical regions from progranulin mutation carriers. (**A**) Human brain tissue from control subjects and patients with FTLD-TDP-*PGRN* was analyzed for granulin peptides and CTSD levels using anti-granulin E and anti-CTSD antibodies. IOC = inferior occipital cortex, MFG = middle frontal gyrus. (**B-D**) Actin-corrected mean band intensities were calculated for 33 kDa granulin E peptide (**B**), 48kDa precursor CTSD (**C**), and 30kDa mature CTSD (**D**), and normalized to the IOC region (n = 4 control subjects, n = 4 FTLD-TDP-*PGRN)*. (**E**) CTSD activity measured in human post-mortem brain tissue for control subjects and patients with FTLD-TDP-*PGRN* (n = 4 control subjects, on the left and n = 5 FTLD-TDP-*PGRN*, on the right, IOC = inferior occipital cortex, MFG = middle frontal gyrus). CTSD activity was normalized to no/low neurodegeneration IOC region. Throughout, values shown are mean ± s.e.m., one-way ANOVA with post-hoc Tukey multiple comparisons test (*P<0.05, **P<0.01, ****P<0.0001, ns = not significant). The aspartyl protease inhibitor Pepstatin A was used as a control for protease activity.

**Table 1.** Clinical information for control and FTLD-TDP-*PGRN* subjects.

| Pathological diagnosis | Age at death | Gender | PMI (hours) |
| --- | --- | --- | --- |
| Control |  |  |  |
| 1 | 81 | F | 30.0 |
| 2 | 86 | F | 7.8 |
| 3 | 76 | M | 8.2 |
| 4 | 86 | F | 6.4 |
| FTLD-TDP-A<br>(*PGRN*-/- ) |  |  |  |
| 1 | 74 | M | 31.0 |
| 2 | 66 | F | 7.4 |
| 3 | 68 | M | 14.0 |
| 4 | 64 | M | 7.2 |
| 5 | 73 | F | 21.0 |

The pathological diagnosis, age at death, gender and post-mortem interval (PMI) for all subjects in Figure 5 is listed.

## Discussion

We have previously shown in *C. elegans* that expression of granulin peptides enhances TDP-43 toxicity and prevents its degradation (Salazar et al., 2015). In this study, we sought to understand the mechanism by which granulins exert their effects and determine if they more broadly impacted protein homeostasis. Granulins are produced in an age and stress-dependent manner to consequently impair ASP-3/CTSD activity, negatively impacting cellular protein homeostasis and driving a lysosomal stress response. These effects manifest as an overall decrease in animal fitness.

This study contributes a key new dimension to our understanding of the regulation of lysosomal proteostasis via the identification of granulins as a novel age and stress-produced peptides that reduce aspartyl protease activity *in vitro*. Indeed, granulins are composed of evolutionarily conserved stacked beta hairpins stabilized by disulfide bonds, which are often found in natural protease inhibitors (Luckett et al., 1999). This highly compact and stable structure is thought to confer resistance to denaturation and protection against proteolytic cleavage. A role for granulins in regulating protease maturation has also been demonstrated in plant cysteine proteases that incorporate a granulin domain C-terminal to the catalytic domain, such as RD21 in *A. thaliana* (Yamada et al., 2001). In support of granulins as regulators of protease activity, homozygous progranulin mutation carriers develop a progressive myoclonic epilepsy syndrome that phenocopies loss of function mutations in another lysosomal protease inhibitor, cystatin B (Pennacchio et al., 1996, Smith et al., 2012).

Granulins likely play a *normal physiological role* in regulating protease activation. Given their ability to promote the CLEAR program, granulins may serve as a signal for stress or impaired health that requires regulated checks on aspartyl protease activity, perhaps to limit inflammation. This would be consistent with the role of progranulin in complement-mediated synaptic pruning by microglia (Lui et al., 2016). We speculate that under conditions of progranulin haploinsufficiency, the normal balance between progranulin and granulins becomes skewed towards excessive granulins. In excess, the inhibitory effect of granulins upon ASP-3/CTSD impairs the function of lysosomes; with age, the natural compensatory mechanisms such as the CLEAR program become overwhelmed, resulting in cellular dysfunction. When this occurs in neurons and/or support cells such as microglia, the end result can be neurodegeneration. Because granulins increase with age, it remains possible that accumulation of granulins directly contribute to the proteostatic pressures associated with increasing age. Comprehensive measures of progranulin-to-granulin ratios with age and in progranulin mutation carriers are underway.

Progranulin is a highly conserved protein (Cenik et al., 2012, Kao et al., 2017, Palfree et al., 2015). Our data suggest that granulin regulation of aspartyl protease activity may be conserved from nematodes to humans. Our computational modeling data suggest that granulin domains may bind to aspartyl protease precursors through the propeptide region. One could then posit that unbinding of the granulin and proteolytic removal of the propeptide are coupled regulatory steps in the endolysosomal compartment. The number of granulin domains has increased through phylogeny from one in *Dictyostelium discoideum* and plants, three in nematodes to seven-and-a-half in humans (Palfree et al., 2015, Yamada et al., 2001). It is intriguing to speculate that this expansion in cleavage fragments could lead to regulation of additional proteases. Indeed, the distinct effects of granulin 2 and 3 on protein homeostasis, lysosome function and TDP-43 toxicity (Salazar et al., 2015), as compared to granulin 1, may suggest functional differences between granulins.

Our results establish age-regulated granulins as modulators of lysosomal aspartyl protease activity, and suggest that a *toxic gain of granulin function*, rather than or in addition to simply loss of full-length progranulin, contributes to FTLD disease pathogenesis. This may explain why progranulin loss-of-function mutations are transmitted in an autosomal dominant fashion. The presence of granulins only in the happloinsufficient state could explain why TDP-43 pathology is not seen in the null state (Smith et al., 2012). Several lysosomal proteases that cleave progranulin have recently been identified (Holler et al., 2017, Lee et al., 2017, Zhou et al., 2017b), which helps to validate our findings. This study now prompts several important follow up questions determining the rate and order in which granulins are liberated from progranulin, exploring how pH changes impact the predicted association of granulins with aspartyl proteases and asking if granulins act more generally in other neurodegenerative disorders such as Alzheimer’s disease. In addition, these studies impact efforts directed towards progranulin repletion as a therapeutic target, as care should be taken to measure whether that progranulin is processed into granulins. Preventing progranulin cleavage into granulins represents a rational new therapeutic target in neurodegeneration.

## Acknowledgements

This work was supported by National Institutes of Health R21NS082709 and R01NS095257, the Alzheimer’s Disease Research Center, the Consortium for Frontotemporal Dementia Research, the UCSF School of Medicine Irene Perstein Award, the John Douglas French Alzheimer’s Foundation and the Brain Research Foundation (A.W.K.). We also thank The James and Barbara Knuppe Family Foundation for research support. Additional support was provided by NIH NIGMS 8P41GM103481, Howard Hughes Medical Institute and NINDS Informatics Center for Neurogenetics and Neurogenomics (P30 NS062691) (A.L.B.), NIH (RC1 AG035610), the NINDS Informatics Center for Neurogenetics and Neurogenomics (P30 NS062691) and the John Douglas French Alzheimer’s Foundation (G.C.), the Rainwater Foundation (A.M.C.) and NIH (R01AG046400) (K.A.). We would like to acknowledge Bradley Webb and Diane Barber for advice on enzymatic assays, and Malene Hansen for providing the MAH235 strain. We thank Alissa Nana Li and the UCSF Neurodegenerative Disease Brain Bank for providing human brain tissue samples, which receives funding support from NIH grants P01AG019724 and P50AG023501, the Consortium for Frontotemporal Dementia Research, and the Tau Consortium. We thank Loan Doan, Ellen Sellinger and Arpana Arjun for assistance with strain maintenance. We thank all members of the Kao lab for helpful discussions. Some strains were provided by the CGC, which is funded by NIH Office of Research Infrastructure Programs (P40 OD010440), and the Japan National BioResource Project.

## Author Contributions

A.W.K. and V.J.B. conceived the project. A.M.C. and B.C. assisted with development of subcellular fractionation protocols for Figure 2 and Figure S2. M.O.P., W.A.C. and M.P.J. contributed to computational modeling data for Figure 4. M.V. and K.A. contributed to short-term associative learning assays for Figure 4I. A.R.A. contributed to immunoblots and enzymatic assays for Figure 5. J.A.O., S.C. and A.L.B. performed all mass spectrometry experiments and analysis. F.G. and G.C. contributed the RNAseq data analysis. W.W.S. and B.L.M. provided human brain tissue samples. V.J.B. contributed to all other immunoblots, behavioral data and enzymatic assays. A.W.K. and V.J.B. prepared the manuscript.

## Author Information

M.P.J. serves on the S.A.B., and is a shareholder, of Schrodinger LLC, which develops, licenses, and distributes software used in this work. Correspondence and requests for materials should be addressed to.

## Materials and Methods

### Strains

*C. elegans* strains were cultured at 20 °C according to standard procedures (Brenner, 1974). Some strains were provided by the Mitani Laboratory (National Bioresource Project, Japan) at the Tokyo Women’s Medical University and the Caenorhabditis Genetics Center (CGC) at the University of Minnesota. Strain descriptions are at www.wormbase.org. The N2E control strain was used as the wild-type strain. The *pgrn-1(tm985)* strain has a 347 bp deletion in the *pgrn-1* gene resulting in a null allele (Kao et al., 2011). The following *C. elegans* strains were used in this study:

<u>CF3050</u> *pgrn-1(tm985) I*

<u>AWK33</u> *pgrn-1(tm985) I; rocIs1[Ppgrn-1*+*SignalSequence::granulin1::FLAG::polycistronic mCherry* + *Punc-122::GFP]*

<u>AWK43</u> *pgrn-1(tm985)* I; *rocEx14[Ppgrn-1*+*SignalSequence::granulin2::FLAG::polycistronic mCherry* + *Pmyo-2::GFP]*

<u>AWK107</u> *pgrn-1(tm985)* I; *rocIs5[Ppgrn-1*+*SignalSequence::granulin3::FLAG::polycistronic mCherry* + *Pmyo-2::GFP]*

<u>AWK524</u> *pgrn-1(tm985) I; muIs189[Ppgrn-1::pgrn-1::polycishronic mCherry* + *Podr-1::CFP]* CF3778 *pgrn-1(tm985) I; muIs213[Ppgrn-1::pgrn-1::RFP]*

<u>AWK181</u> *pgrn-1(tm985) I; unc-119(ed3)III; pwIs503[vha6p::mans::GFP* + *Cb unc-119(*+*)]; muIs213[Ppgrn-1::pgrn-1::RFP]*

<u>AWK360</u> *pgrn-1 (tm985) I; unc-119(ed3) III; pwIs50[Plmp-1::lmp-1::GFP* + *Cbr-unc-119(*+*)]; muIs213[Ppgrn-1::pgrn-1::RFP]*

<u>AWK395</u> *pgrn-1 (tm985) I; unc-119(ed3) III; cdIs54[pcc1::MANS::GFP* + *unc-119(*+) + *myo-2::GFP]; muIs213[Ppgrn-1::pgrn-1::RFP]*

<u>AWK374</u> *pgrn-1 (tm985) I; bIs34[rme-8::GFP* + *rol-6(su1006)]; muIs213[Ppgrn-1::pgrn-1::RFP]*

<u>DA2123</u> *adIs2122 [lgg-1p::GFP::lgg-1* + *rol-6(su1006)]*

<u>AWK393</u> *pgrn-1 (tm985) I; adIs2122[lgg-1p::GFP::lgg-1* + *rol-6(su1006)]*

<u>AWK432</u> *pgrn-1 (tm985) I; adIs2122[lgg-1p::GFP::lgg-1* + *rol-6(su1006)]; rocIs1[Ppgrn-1*+*SS::granulin1::FLAG::polycistronic mCherry]*

<u>AWK434</u> *pgrn-1 (tm985) I; adIs2122[lgg-1p::GFP::lgg-1* + *rol-6(su1006)]; rocEx14[Ppgrn-1*+*SS::granulin2::FLAG::polycistronic mCherry* + *Pmyo-2::GFP]*

AWK435 *pgrn-1 (tm985) I; adIs2122[lgg-1p::GFP::lgg-1* + *rol-6(su1006)]; rocIs5[Ppgrn-1*+*SS::granulin3::FLAG::polycistronic mCherry* + *Pmyo-2::GFP]*

<u>AWK436</u> *pgrn-1 (tm985) I; muIs189[Ppgrn-1::pgrn-1::polycishronic mCherry* + *Podr-1::CFP]; adIs2122[lgg-1p::GFP::lgg-1* + *rol-6(su1006)]*

<u>MAH235</u> *sqIs19[Phlh-30::hlh-30::gfp* + *rol-6(su1006)]*

<u>AWK403</u> *pgrn-1(tm985) I; sqIs19[Phlh-30::hlh-30::gfp* + *rol-6(su1006)]*

<u>AWK404</u> *pgrn-1(tm985) I; sqIs19[Phlh-30::hlh-30::gfp* + *rol-6(su1006)]; rocIs1[Ppgrn-1*+*SS::granulin1::FLAG::polycistronic mCherry]*

<u>AWK405</u> *pgrn-1(tm985) I; sqIs19[Phlh-30::hlh-30::gfp* + *rol-6(su1006)]; rocEx14[Ppgrn-1*+*SS::granulin2::FLAG::polycistronic mCherry* + *Pmyo-2::GFP]*

<u>AWK406</u> *pgrn-1(tm985) I; sqIs19[Phlh-30::hlh-30::gfp* + *rol-6(su1006)]; rocIs5[Ppgrn-1*+*SS::granulin3::FLAG::polycistronic mCherry* + *Pmyo-2::GFP]*

<u>JIN1375</u> *hlh-30(tm1978) IV*

<u>AWK514</u> *pgrn-1 (tm985) I; hlh-30(tm1978) IV*

<u>AWK516</u> *pgrn-1 (tm985) I; hlh-30(tm1978) IV; rocIs1[Ppgrn-1*+*SS::granulin1::FLAG::polycis mCherry]*

<u>AWK518</u> *pgrn-1 (tm985) I; hlh-30(tm1978) IV; rocEx14 [Ppgrn-1*+*SS::granulin2::FLAG::polycistronic mCherry* + *Pmyo-2::GFP]*

<u>AWK519</u> *pgrn-1 (tm985) I; hlh-30(tm1978) IV; rocIs5 [Ppgrn-1*+*SS::granulin3::FLAG::polycistronic mCherry* + *Pmyo-2::GFP]*

<u>AWK521</u> *pgrn-1 (tm985) I; hlh-30(tm1978) IV; muIs189[Ppgrn-1::pgrn-1::polycistronic mCherry* + *Podr-1::CFP]*

<u>AWK296</u> N2E; *Ex[Pced-1::asp-3::mrfp* + *pRF4(rol-6)]; unc-119(ed3) III; pwIs50[Plmp-1::lmp-1::GFP* + *Cbr-unc-119(*+)]

<u>AWK333</u> *pgrn-1(tm985) I; Ex[Pced-1::asp-3::mrfp* + *pRF4(rol-6)]; unc-119(ed3) III; pwIs50[Plmp-1::lmp-1::GFP* + *Cbr-unc-119(*+)]

<u>AWK334</u> *pgrn-1(tm985) I; Ex[Pced-1::asp-3::mrfp* + *pRF4(rol-6)]; unc-119(ed3) III; pwIs50[Plmp-1::lmp-1::GFP* + *Cbr-unc-119(*+*)]; rocIs5[Ppgrn-1*+*SS::granulin3::FLAG::polycis tronic mCherry* + *Pmyo-2::GFP]*

<u>AWK177</u> *asp-3(tm4450) X*

<u>TU38</u> *deg-1(u38) X*

<u>CF3884</u> *pgrn-1(tm985) I; deg-1(u38) X*

<u>AWK224</u> *pgrn-1(tm985) I; deg-1(u38) X; rocIs5[Ppgrn-1*+*SS::granulin3::FLAG::polycis tronic mCherry* + *Pmyo-2::GFP]*

<u>VM487</u> *nmr-1(ak4)II*

### Generation of transgenic *C. elegans*

To generate strains expressing individual granulins, each granulin was amplified separately from wild-type *C. elegans* progranulin cDNA as previously described (Salazar et al., 2015).

### ER stress assays

ER stress assays were performed as previously described (Judy et al., 2013).

### Animal viability

L4 stage animals were allowed to lay eggs overnight. Fifty synchronized eggs were transferred to seeded plates. After three days, the fraction of animals that developed to the L4 stage was quantified.

### Body size

L4 animals were picked, grown at 20 °C overnight and imaged the following day as day 1 adults. Animals were mounted on a 2 % agarose pad with 25 mM sodium azide (Spectrum Chemical, #SO110) and imaged using a Zeiss AxioImager microscope at 10x. Body size was measured in ImageJ software using the skeletonize function.

### Short-term associative learning

Short-term associative learning assays were performed as previously described (Kauffman et al., 2010, Torayama et al., 2007).

### Immunoblotting for progranulin expression and cleavage

Sixty L4 stage animals were allowed to lay eggs overnight (~sixteen hours). Adult worms and hatched larvae were washed off the plates with M9 buffer. Eggs were collected with a cell scraper and transferred to a newly seeded plate by chunking. These eggs were allowed to develop to early L4 stage and 200 μl 20 mM FUDR (Fisher Scientific, #AC227601000) was added to prevent development of progeny and overgrowth of plates. At the appropriate time point, animals were collected from plates with ice cold M9 and washed once to remove food. The worm pellet was resuspended 1:1 in freshly made ice cold RIPA buffer (50 mM Tris pH 7.4, 150 mM NaCl, 5 mM EDTA, 0.5 % SDS, 0.5 % SDO, 1 % NP-40, 1 mM PMSF, cOmplete protease inhibitor (Roche, #04693124001) and PhosSTOP phosphatase inhibitor (Roche, #04906837001), 0.3 mM Pefabloc (Roche, #11429868001). Worms were transferred to Eppendorf tubes and sonicated for 4 cycles of 1 minute on and 2 minutes off (BioRuptor, Diagenode). Lysates were centrifuge for 5 minutes at 13,000 rpm at 4 °C. Supernatant was transferred to a fresh Eppendorf tube and samples were boiled at 95 °C (with 4x LDS, 10 % reducing agent) for 5 minutes and analyzed by SDS PAGE. 50 μg total protein was resolved on 4-12 % gradient SDS-PAGE gels and transferred to PVDF.

### Antibodies

Commercial antibodies used for Western blotting were the following:

Anti-HSP-4/BiP (Novus Biologicals, #NBP1-06274, 1:1000 dilution)

Anti-RFP (GenScript, #A00682, 1:1000 dilution)

Anti-GFP (ThermoFisher, #MA5-15256, 1:1000 dilution)

Anti-FLAG (Sigma, #F3165, 1:1000 dilution)

Anti-cathepsin D (R&D Systems Inc., #AF1014, 1:200 dilution)

Anti-HSP-70/HSC-70 (Santa Cruz Biotechnology Inc., #sc-33575, 1:1000 dilution)

Anti-calnexin (Novus Biologicals, #NBP1-97476, 1:1000 dilution)

Anti-LMP-1(Developmental Studies Hybridoma Bank, #LMP1, 1:100 dilution)

Anti-actin (EMD Millipore, #MAB1501R, 1:5000 dilution)

Anti-tubulin (Abcam, #ab50721, 1:1000 dilution)

Goat anti-mouse (LI-COR IRDye 800CW, #925-32210, 1:10,000 dilution)

Goat anti-rabbit (LI-COR IRDye 800CW, #925-32211, 1:10,000 dilution)

Donkey anti-goat (LI-COR IRDye 800CW, #925-32214, 1:10,000 dilution)

Donkey anti-mouse (LI-COR IRDye 680RD, #925-68072, 1:10,000 dilution)

Antibodies made in-house and used for Western blotting were the following:

Anti-granulin 1 (RB2481, Biomatik, epitope HQCDAETEC(acm)SDDET, 1:1000 dilution) Anti-granulin 2 (RB2483, Biomatik, epitope CPDKASKC(acm)PDGST, 1:1000 dilution) Anti-granulin 3 (RB2487, Biomatik, epitope CTVLMVESARSTLKL, 1:1000 dilution) Anti-granulin E (RB4367, Biomatik, epitope ECGEGHFCHDNQTCCR, 1:100 dilution)

Imaging and quantification were performed on the LI-COR Odyssey Infrared System. Three independent blots were performed.

### Confocal microscopy

Animals were mounted on microscope slides with 2 % agarose pads containing 30 mM levamisole hydrochloride (Fisher Scientific, #AC187870100) and imaged using a Zeiss LSM 700 laser-scanning confocal microscope using 488 nm and 561 nm lasers and 63x and 100x objectives. L1 animals were imaged 1-2 h after hatching. Z-stacks were taken every 0.7 μm. Image processing was carried out using ImageJ software. A maximum intensity projection of the z-stack for each animal was created. Images at 488 nm and 561 nm were overlaid and analyzed for co-localization.

### Subcellular fractionation

Thirty L4 stage animals were picked to 50 x 10 cm plates per strain. Plates were confluent with mixed stage animals after four days growth at 20 °C. Progranulin cleavage was observed after starving animals for an additional seventy-two hours at 20 °C. A lysosome fraction was isolated from a light mitochondrial-lysosome fraction as previously described (Cuervo et al., 1997) with the following modifications. Animals were collected in 0.25 M sucrose (pH 7.2) and washed twice with 0.25 M sucrose. Lysosomes and mitochondria were separated using a discontinuous Nycodenz (Progen Biotechnik, Germany, #1002424) density gradient. Lysosomes were collected from the 19.8 % / sucrose interface and the 26.3 / 19.8 % interface and pooled. Lysosomes were diluted five times with 0.25 M sucrose, and pelleted at 37,000 × *g* for 15 minutes. Cytosolic, ER and lysosomal fractions were confirmed by immunoblotting for specific subcellular fraction markers (LAMP-1, HSC-70, calnexin).

### Co-immunoprecipitation of FLAG-tagged granulin 3 and mass spectrometry analysis

Granulin 3 expressing animals were grown on 10 x 15 cm high growth medium plates, collected and washed in M9 buffer, and re-suspended in lysis buffer with protease inhibitor (20 mM HEPES, 150 mM NaCl, 2 mM EDTA, 0.1 % Triton-X, 10 % glycerol, 40 mM NaF, cOmplete protease inhibitor (Roche, #04693124001). Worms were drop frozen in liquid nitrogen and lysed by grinding using a mortar and pestle. The FLAG-tagged granulin protein was captured using a FLAG-immunoprecipitation kit (Sigma-Aldrich, #FLAGIPT1). Bound proteins were eluted and separated by SDS-PAGE, silver stained and digested with trypsin. Peptides recovered were analyzed by reversed-phase liquid chromatography-electrospray tandem mass spectrometry (LC-MS/MS) in a hybrid linear ion trap-Orbitrap mass spectrometer (LTQ-OrbitrapVelos, ThermoScientific, San Jose, CA). Data generated from spectra were searched against the *C. elegans* subset of the UniProtKB database 06 17 2013 using in-house ProteinProspector (Clauser et al., 1999). For each protein identified, a corrected abundance index was calculated (peptide counts / molecular weight of the protein). The corrected abundance of each protein in the control (progranulin null) and experimental (granulin-expressing) lane were plotted against each other and the line of best fit determined. The standard deviation of all points to the line of best fit was calculated and interaction candidates were identified above three standard deviations distance from the line of best fit. Additional candidates were identified as being pulled down only in the experimental lane and not in the control lane.

### ASP-3/CTSD activity measurements from total worm lysates

ASP-3/CTSD activity was measured using a commercially available kit (BioVision Cathepsin D Activity Fluorometric Assay Kit, #K143-100). Animals were staged as for immunoblotting, but without the addition of 20 mM FUDR. At day 1 adulthood, worms were collected from plates with ice cold M9 and washed twice to remove food. Worm pellets were resuspended in 1 x RIPA buffer (Fisher Scientific) and frozen at −80 °C overnight. Pellets were thawed and sonicated for 4 cycles of 1 min on and 2 min off (BioRuptor, Diagenode). Lysates were centrifuged for 5 minutes at 13,000 rpm at 4 °C and supernatant was transferred to a fresh tube. 0.25 μg total protein per sample was used per assay and samples from one strain were run in triplicate. Fluorescence measurements were taken every minute at 25 °C (Infinite M200, Tecan). For the negative control, 250 nM Pepstatin A was added to the lysate and pre-incubated for 10 minutes on the bench at room temperature. Linear regression was performed on at least 30 minutes of data to calculate the rate of enzyme activity.

### Necrotic cell counts

Animals were anesthetized on 2% agarose pads containing 30 mM levamisole hydrochloride (Fisher Scientific, #AC187870100) mounted onto glass slides and imaged using a Zeiss AxioImager microscope with 5X and 63X DIC objectives. Necrotic PVC interneurons were counted at 63X magnification in 100 animals at L2, L3, L4, and Day 1 adulthood. Each developmental time point per strain was assayed three independent times.

### Homology modeling of *C. elegans* granulin domains

Models of *C. elegans* granulin domains 1, 2, and 3 were generated by homology modeling using the Structure Prediction Wizard module of Schrödinger’s Prime program (Schrödinger, 2015, Jacobson et al., 2002, Jacobson et al., 2004). The FASTA sequences of *C. elegans* granulins were obtained from UniProt (UniProt, 2015). Using the NMR solution structure of human granulin A as a template (PDB ID: 2JYE) (Tolkatchev et al., 2008), a homology model was built for the *C. elegans* granulin domains. The homology models were then energy-refined using the Refine Loops panel of the Prime program (Schrödinger, 2015). In order to further refine the characteristic stacked beta hairpin conformation of granulin, 20 ns of restrained molecular dynamics (MD) simulation was carried out for each homology model, followed by 10 ns of unrestrained MD simulation, using the AMBER 14 suite of programs (D.A. Case, 2015). Each system was solvated with TIP3P water (William L. Jorgensen, 1983) and counterions were added to neutralize the system by tleap program (Schafmeister, 1995). Water was first minimized and simulated for 150 ps in the NPT ensemble with a harmonic restraint of 2.0 kcal/mol Å^2^ on the protein to relax the water. The entire system was then minimized and heated to 300 K over 500 ps. Two equilibrations with respective duration of 200 ps were performed using harmonic distance, angle, and dihedral restraints of 20 kcal/mol Å^2^ on the protein backbone based on the NOE contacts reported for carp granulin 1 (Hrabal et al., 1996). First, the system was equilibrated at constant volume and temperature (NVT) using a Langevin thermostat (Uberuaga et al., 2004). Following this, the second equilibration was performed at constant pressure and temperature (NPT) using a Berendsen barostat (Yuhong Zhang, 1995) with isotropic position scaling to bring the system to a stable density. A 20 ns production was then performed in the NVT ensemble with the restraints, followed by a 10 ns unrestrained simulation to relax the system. The Particle Mesh Ewald summation method was used to compute long-range electrostatic interactions (Tom Darden, 1993, Ulrich Essmann, 1995), and short-range nonbonded interactions were truncated at 8 Å in the periodic boundary conditions. All dynamics are carried out using the *pmemd.cuda* module of AMBER 14 suite of programs (D.A. Case, 2015, Uberuaga et al., 2004, Gotz et al., 2012).

### Homology modeling of *C. elegans* ASP-3 protease and protein-protein docking

The FASTA sequence of the *C. elegans* protease ASP-3 were obtained from UniProt (UniProt, 2015). The precursor of the aspartyl protease ASP-3 was modeled with the target-template alignment using ProMod3 and Swiss-Model (Guex et al., 2009) based on the X-ray crystallographic structure of porcine pepsinogen (PDB ID: 2PSG). The Z-DOPE score was used to assess the quality of the model, with the precursor of ASP-3 yielding a score lower than −1.0, indicative of acceptable quality models (Shen and Sali, 2006). Protein-protein docking was then carried out between *C. elegans* proASP-3 and granulins 1, 2 and 3 using the ZDock server (Pierce et al., 2014). The top 10 poses with the lowest energy values were chosen for a prediction of the binding site of granulins to proASP-3.

### HLH-30/TFEB imaging

Forty L4 animals were picked, grown at 20 °C overnight and imaged the following day as day 1 adults. The nuclear localization of HLH-30::GFP was imaged using a Zeiss AxioImager microscope at 10x. Animals were imaged within 5 minutes of mounting on a 2 *%* agarose pad with 25mM sodium azide (Spectrum Chemical, #SO110). Data from three independent experiments were pooled.

### RNA sequencing analysis

Total RNA was isolated from wild-type (N2E), *pgrn-1(tm985)*, and *pgrn-1(tm985); granulin 1(+)* expressing animals using chloroform extraction, and DNA contamination was removed with DNA-free treatment (Ambion). Samples were extracted in quadruplicate (four biological replicates for each strain), for a total of 12 samples. Total RNA was quantified using the RiboGreen assay (ThermoFisher, #R11490) and RNA quality was checked using an Agilent Bioanalyzer (Agilent). RNA Integrity Numbers (RINs) were >8 in all the samples. Libraries for RNA-seq were prepared using the Illumina TruSeq library preparation protocol (Illumina Inc), multiplexed into a single pool and sequenced using an Illumina HiSeq 2500 sequencer across 4 lanes of 2 Rapid Run SR 1650 flow cells. After demultiplexing, we obtained between 13 and 32 million reads per sample, each one 50 bases long. Quality control was performed on base qualities and nucleotide composition of sequences. Alignment to the *C. elegans* genome (ce10) was performed using the STAR spliced read aligner (Dobin et al., 2013) with default parameters. Additional QC was performed after the alignment to examine the following: level of mismatch rate, mapping rate to the whole genome, repeats, chromosomes, and key transcriptomic regions (exons, introns, UTRs, genes). Between 92 and 93 % of the reads mapped uniquely to the worm genome. Total counts of read fragments aligned to candidate gene regions within the *C. elegans* reference gene annotation were derived using HTS-seq program and used as a basis for the quantification of gene expression. Only uniquely mapped reads were used for subsequent analyses. Following alignment and read quantification, we performed quality control using a variety of indices, including sample clustering, consistency of replicates, and average gene coverage. Differential expression analysis was performed using the EdgeR Bioconductor package (Robinson et al., 2010), and differentially expressed genes were selected based on False Discovery Rate (FDR, Benjamini-Hochberg adjusted p-values) estimated at ≤ 5 %. Clustering and overlap analyses were performed using Bioconductor packages within the statistical environment R (www.rproject.org/). Gene Ontology annotation was performed using DAVID (david.abcc. ncifcrf.gov/).

### TFEB binding site analysis

The promoter regions of all differentially regulated transcripts were analyzed for the presence of the *C. elegans* TFEB/HLH-30 binding site E-box sequence 5’-CACGTG-3’. Enrichment of TFEB binding sites was tested by comparison to the expected distribution based on 1000 random permutations. A two-sided Student’s t-test was used to calculate p-values.

### CTSD activity measurements from human brain tissue

Brain tissue was prepared as previously described (Salazar et al., 2015). Briefly, frozen human brain tissue was obtained from the UCSF Neurodegenerative Disease Brain Bank. Subjects were chosen based on a postmortem interval < 48 h (range 6.4–31.0 h). Adjacent tissue blocks were fixed, embedded in paraffin wax, sectioned, stained for hematoxylin and eosin, and rated for astrogliosis (0–3 scale). Frozen tissue blocks were chosen for inclusion based on a level of severe (3 high gliosis) or absent (0–1 low gliosis) neurodegeneration in the adjacent fixed tissue block. Using these criteria, we selected the middle frontal gyrus (severe neurodegeneration) and inferior occipital cortex (absent to low neurodegeneration) to measure cathepsin activity. Brain tissue samples were weighed, diluted 25-fold with RIPA buffer (Alfa Aesar, #J62524), homogenized for 1 minute (pestle pellet motor) and sonicated for 5 rounds of 30 seconds on and 1 minute off (BioRuptor, Diagenode). 0.25 μg total protein per sample was used per assay and samples were run in triplicate. Fluorescence measurements and analysis were performed as described above for total worm lysates but at 37 °C.

### Data availability

The data that support the findings of this study are available from the corresponding author upon reasonable request.

## Supplementary Figures

**Supplementary Figure 1.**
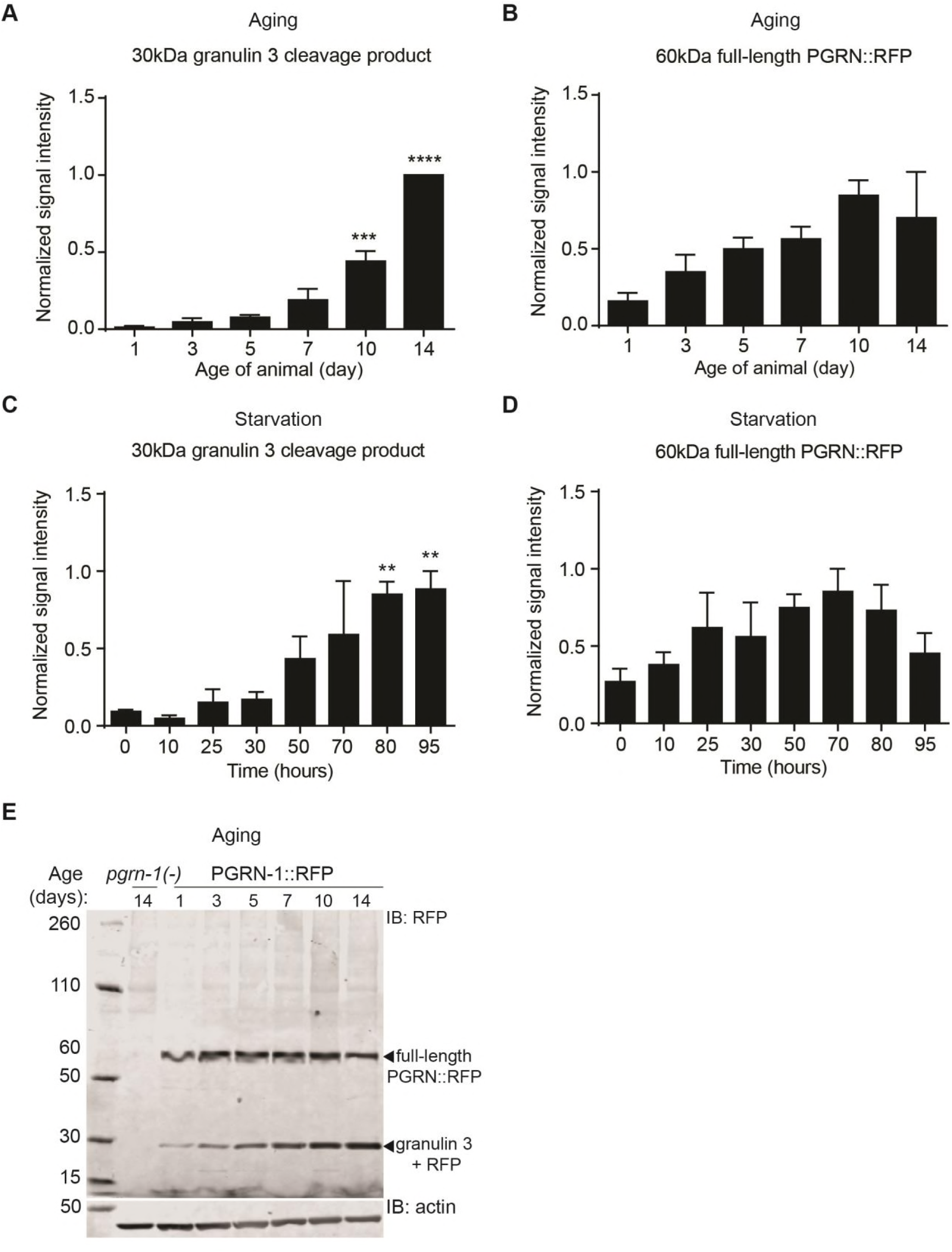
Progranulin expression and cleavage is stress-responsive. (**A-D**) Quantification of actin-corrected mean band intensities from Western blot of *C. elegans* PGRN-1::RFP lysates with aging (**A-B**), and starvation (**C-D**). Actin-corrected mean band intensities were normalized to highest value per experiment (data from three biological replicates is shown, values shown are mean ± s.e.m., one-way ANOVA and Tukey multiple comparisons test, **P<0.01, ***P<0.001, ****P<0.0001). (**E**) In *C. elegans* PGRN-1::RFP lysates the most prominent cleavage product at ~30kDa was recognized by both granulin 3 and RFP antibodies.

**Supplementary Figure 2.**
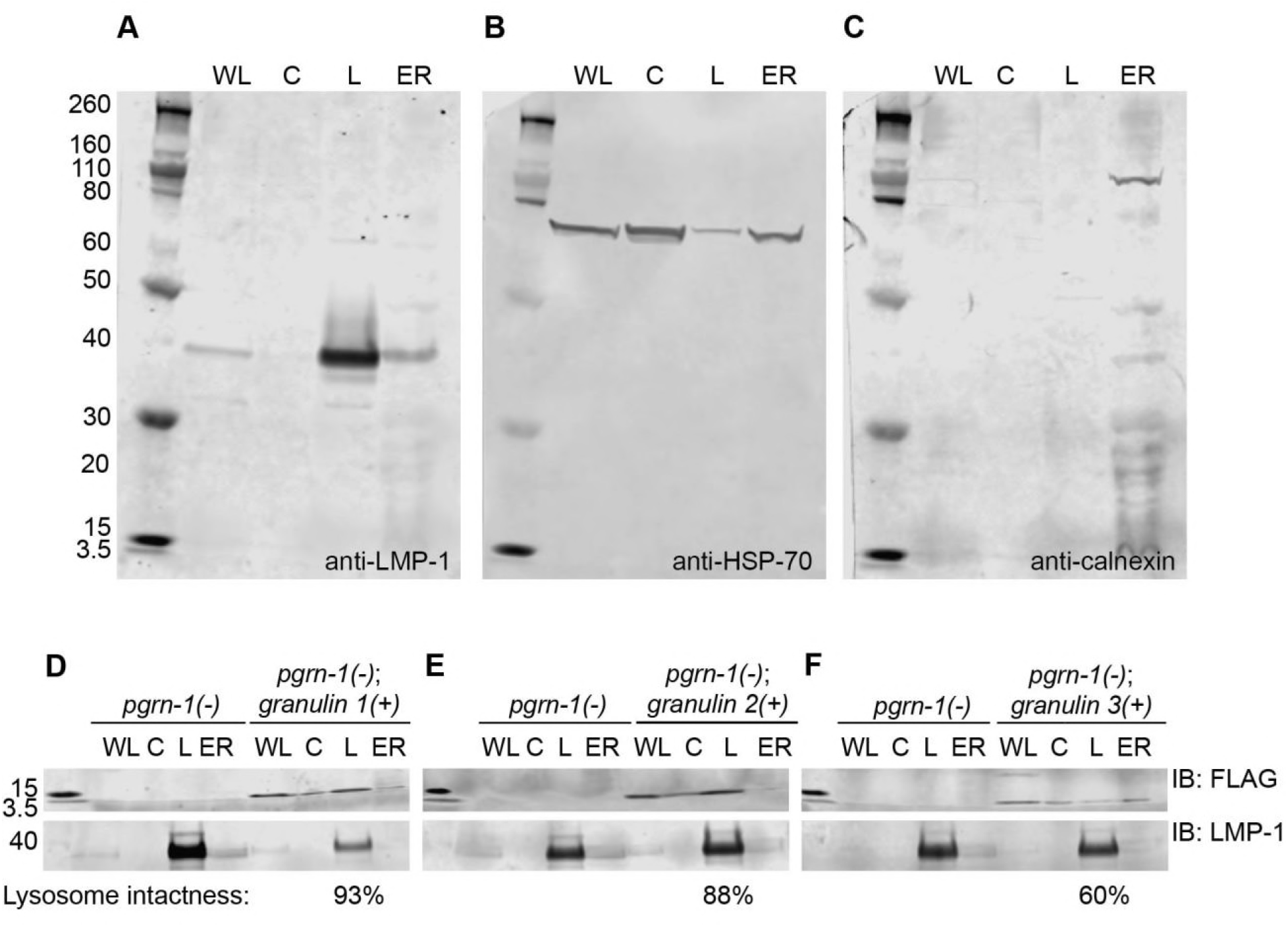
Expressed granulins are found within lysosomes. (**A-C**) Validation of subcellular fractions from *C. elegans* by blotting for fraction-specific markers: anti-LMP-1 is specific for the lysosomal fraction (**A**), anti-HSP-70 which is normally found in both cytosolic and lysosome fractions (**B**), and calnexin which localizes to the ER fraction (**C**). Similar results were observed in three independent Western blots. WL = whole lysate, C = cytosol, L = lysosomes, ER = endoplasmic reticulum. (**D**) Granulin 1, (**E**) granulin 2 and (**F**) granulin 3 localize in the lysosomal fraction. 10 μg of total protein was loaded into each fraction and Western blots were probed with anti-FLAG and anti-LMP-1 antibodies.

**Supplementary Figure 3.**
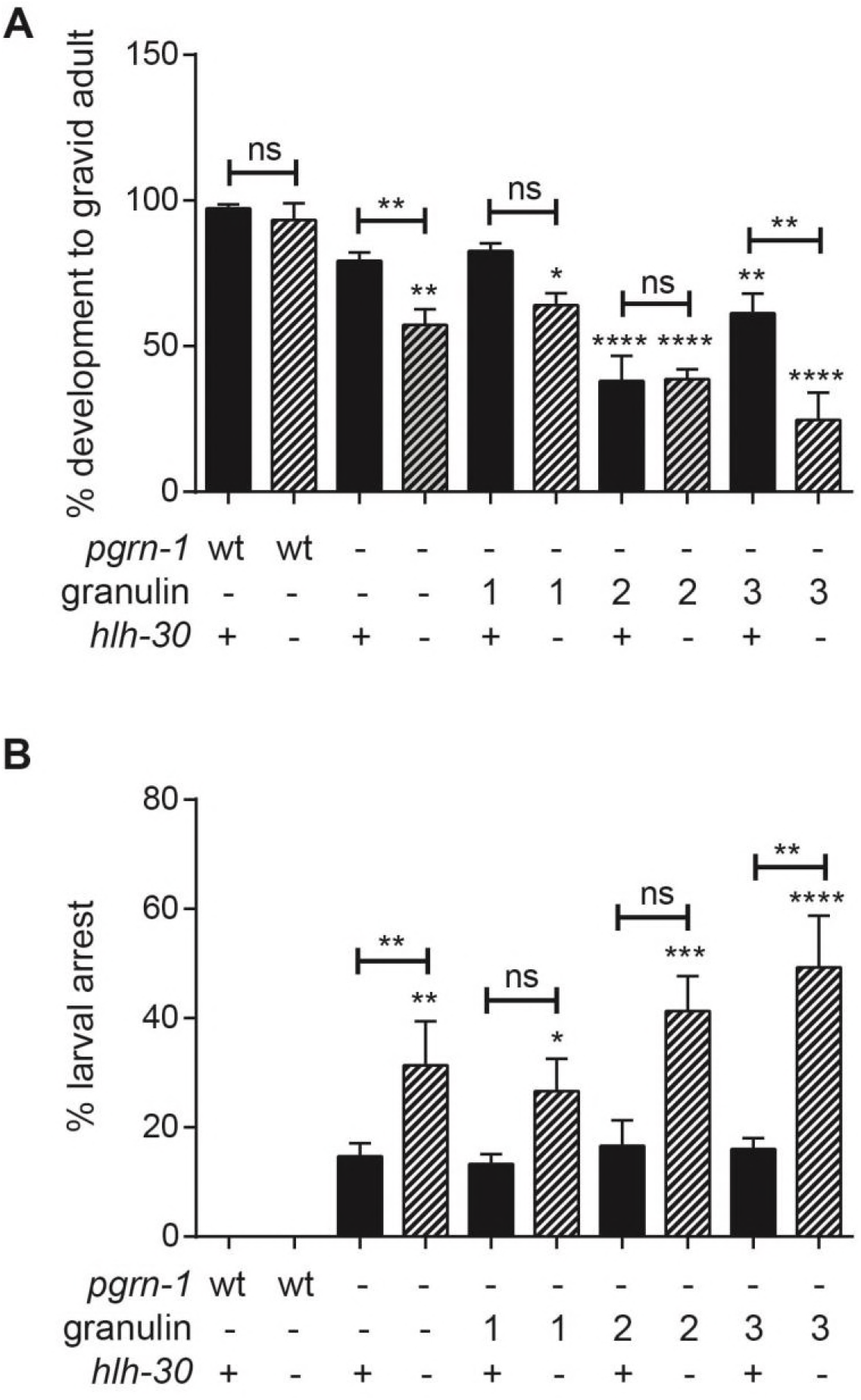
The granulin-induced CLEAR response is a compensatory transcriptional response and requires *hlh-30*. Wild-type, *pgrn-1(-)* and granulin-expressing animals with and without *hlh-30* expression were staged as embryos, and animals were scored for (**A**) development to gravid adult (n = 50, 3 biological replicates), and (**B**) the number of larvae arresting at L1 and L2 stage (n = 50, 3 biological replicates). Values shown are mean ± s.e.m., one-way ANOVA and Tukey multiple comparisons test. Comparisons are to wild-type unless otherwise indicated (*P<0.05, **P<0.01, ***P<0.001, ****P<0.0001, ns = not significant).

**Supplementary Table 1.** Identification of transcripts differentially regulated on progranulin loss and granulin expression. (See corresponding Excel file).

**Supplementary Table 2.** Identification of TFEB binding sites in differentially regulated transcripts from progranulin loss and granulin expression. (See corresponding Excel file).

**Supplementary Table 3.**
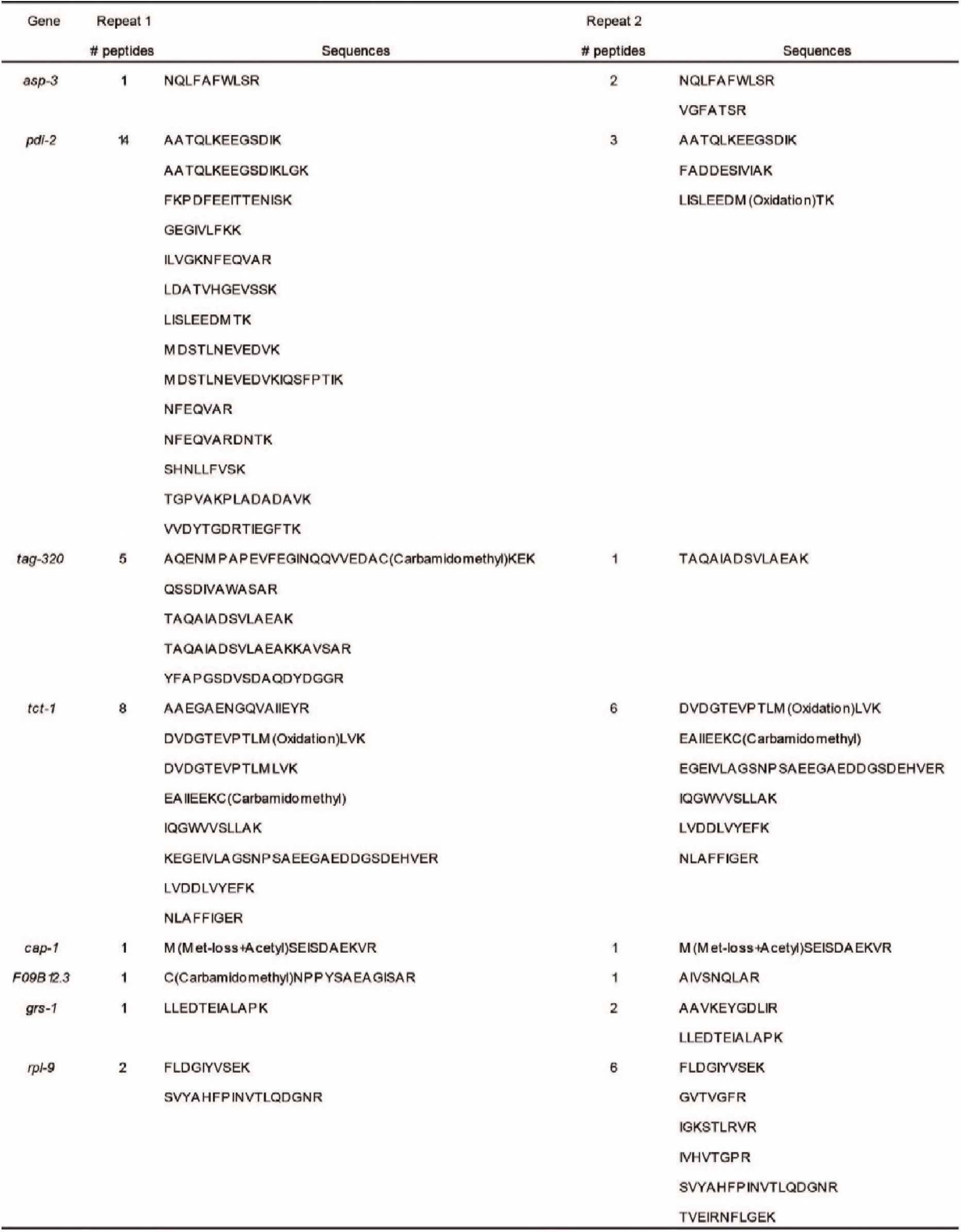
Candidate interactors identified by co-immunoprecipitation using *C. elegans* granulin3::FLAG.

